# Global integrators of light signalling and nutrient homeostasis safeguard symbiosis under light stress

**DOI:** 10.64898/2026.07.29.741514

**Authors:** Moritz Sexauer, Maria-Fotini Anastasakis, Michael Riedel, Samuel B. Blaum, Jana Reichert, Michael Stollenwerk, Ann-Kristin Lenz, Katharina Markmann

## Abstract

Their large, protein-rich seeds render many legume plants like beans, peas or soy attractive food or fodder crops. This is thanks to nitrogen-fixing root nodulation symbiosis (RNS) with rhizobial bacteria, a mutualistic association enabling legumes to not only form storage seeds, but also thrive in nitrogen poor habitats. These benefits come at a cost: accommodating millions of bacteria that actively fix N_2_ requires substantial nutrient and energy input. Evolutionarily successful hosts employ a multi-layered regulatory system balancing endogenous nutrient status and interorganismal nutrient exchange with symbiosis establishment and progression. Root infection as well as nodule organogenesis and –lifespan are tightly regulated, ensuring a mutualistic status. Under adverse conditions, established nodules can be deactivated by induction of nodule senescence in a process involving the transcription factor genes *NAM ATAF CUC* (*NAC*) *094* (Wang et al., 2023) and *FIXATION UNDER NITRATE* (*FUN*) *(Lin et al., 2024).* The molecular basis and role of senescence induction in RNS regulation is still far from understood. Here, we demonstrate that in *Lotus japonicus*, the two photomorphogenesis genes *ELONGATED HYPOCOTYL5* (*HY5*) and the B-Box-containing zinc finger transcription factor gene *BBX21* are required and sufficient for maintaining nodule function under high light intensities. Loss of either *HY5* or *BBX21* results in a light-dependent increase of *NAC094* transcript levels, nodule necrosis and early senescence, and enhanced nodule proliferation. Moreover, expression of dominant negative *HY5* or *BBX21* is sufficient to induce nodule necrosis and hypernodulation. Our data establish *LjHY5* and *LjBBX21* as a novel gene pair safeguarding nodule function and nitrogen fixation by preventing light induced senescence.

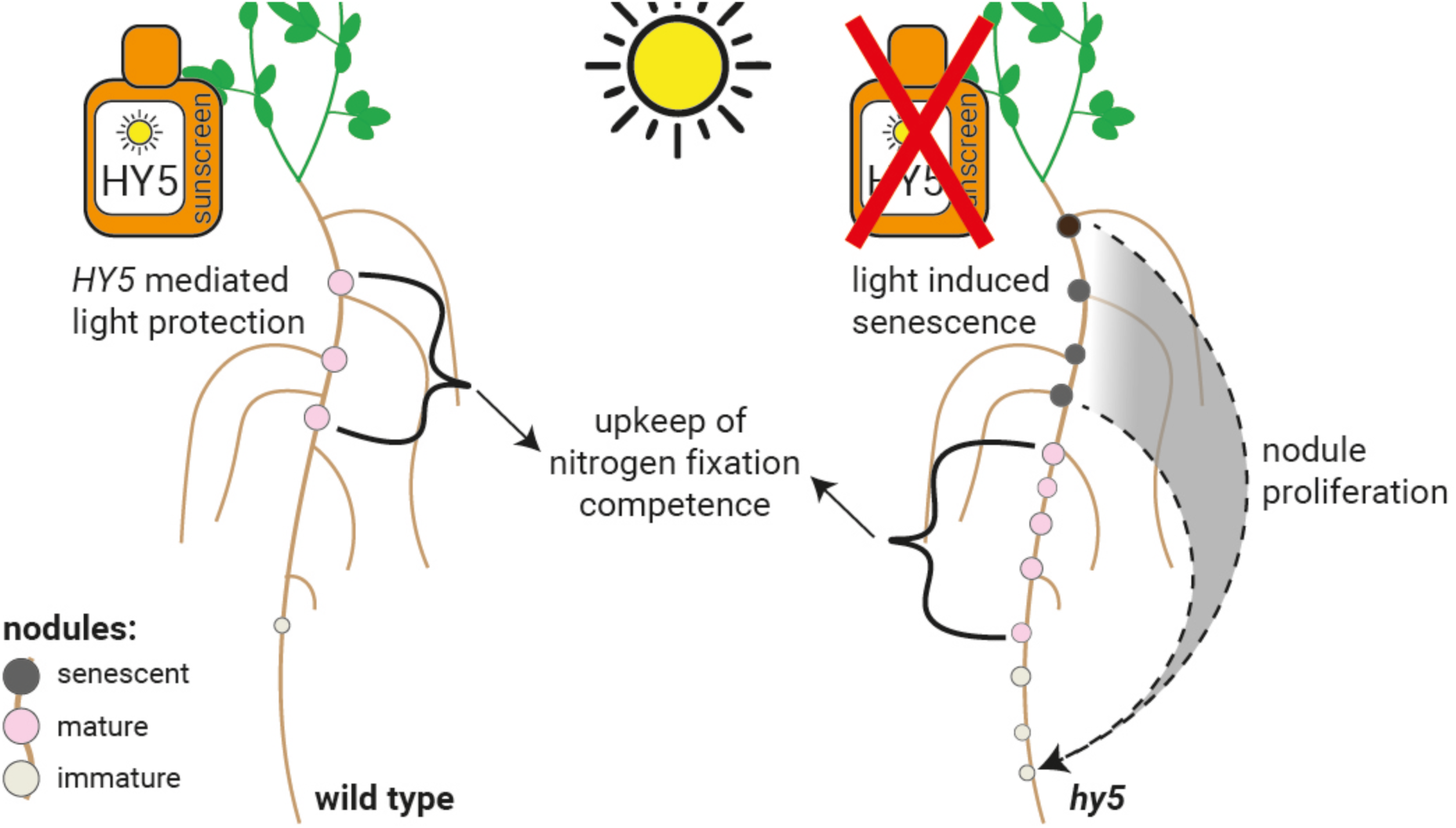
Graphical abstract.

## Introduction

The great majority of plants are photosynthetic carbon autotrophs able to produce sugars *via* carbon dioxide fixation. Besides carbon, organic matter contains predominantly hydrogen, oxygen and nitrogen, with the latter constituting a growth limiting factor in many terrestrial ecosystems (LeBauer and Treseder, 2008). Similarly, plants are nitrogen autotrophic. Yet, they cannot utilize atmospheric dinitrogen (N_2_). Thus, the availability of accessible nitrogen compounds in the rhizosphere is vital for plant growth. Most legumes overcome this constraint by establishing nitrogen fixing root nodule symbiosis (RNS) with rhizobial bacteria. Rhizobia are accommodated in specialized root organs called nodules. These represent sites of intense nutrient exchange, where hosts supply rhizobia with photoassimilates as well as macro– and micronutrients in exchange for ammonia (Roy et al., 2019). Tight regulation of symbiosis by a plant genetic network called Autoregulation of Nodulation (AON) maintains the interaction at a mutualistic status. The host thereby ensures a fitness gain by preventing bacterial parasitism (Li et al., 2022). AON systemically controls infection and nodule establishment *via* systemic signals navigating either xylem vessels in a root-shoot directionality, or phloem sap moving from shoot source tissues to the root (Valmas et al., 2023). These signals comprise components maintaining symbiotic susceptibility, such as CEP peptides and miR2111 (Tsikou et al., 2018; Gautrat et al., 2020), as well as negative regulators of RNS including CLE peptides (Okamoto et al., 2013; Nishida and Suzaki, 2018). Beyond bacterial infection and nodule organogenesis, plants adjust nodule function to their nitrogen demand. In line with this, nitrate supply triggers nodule senescence (Nishida and Suzaki, 2018). This response depends on *NIN LIKE PROTEIN* (*NLP)1/4* and *NITRATE TRANSPORTER (NRT)2.1* mediated nitrate signaling, *FIXATION UNDER NITRATE* (*FUN*) and the NAC-type transcription factor *NAC094* (Wang et al., 2023; Lin et al., 2024).

Light enables photosynthesis. Consistently, light perception by conserved sensors such as *PHYTOCHROMEs (PHYs)* and *CRYPTOCHROMEs* (*CRYs*) induces signaling responses that inform cells and tissues of photosynthate availability. Simultaneously, they trigger appropriate developmental processes jointly referred to as photomorphogenesis (Fankhauser and Chory, 1999). Consistent with nodules representing strong sink tissues, light signaling was shown to regulate nodule number via *PHYs* and *CRYs* (Suzuki et al., 2011; Shimomura et al., 2016; Wang et al., 2021). Light-induced regulation of nodulation involves homologues of *LONG HYPOCOTYL 5* (*HY5*) (Nishimura et al., 2002a; Wang et al., 2021; Ji et al., 2022), a highly promiscuous transcription factor playing a key role in mediating light signaling in Arabidopsis (Osterlund et al., 2000; Lee et al., 2007; Xiao et al., 2021). *HY5* is functionally conserved across angiosperm lineages. It was shown to regulate photomorphogenesis, shade avoidance, and light-induced flavonoid production not only in Arabidopsis, but similarly in tomato, maize and grape (Xiao et al., 2021). Targeting ACGT-containing cis-elements, HY5 is predicted to enhance or repress over 3000 different loci in Arabidopsis (Datta et al., 2007; Lee et al., 2007; Xiao et al., 2021). In darkness, CONSTITUTIVE PHOTOMORPHOGENIC 1 (COP1) activity leads to HY5 ubiquitination and degradation in Arabidopsis (Hardtke et al., 2000; Osterlund et al., 2000; Xu, 2020). Light signaling suppresses COP1 activity, resulting in HY5 protein accumulation (Podolec and Ulm, 2018; Han et al., 2020). HY5 protein activity is modulated by a group of B-BOX (BBX) proteins, which either enhance or suppress HY5 functionality by recruiting it into hetero-dimers (Gangappa et al., 2013; Lin et al., 2018; Bursch et al., 2020; Xu, 2020; Xiao et al., 2021). For example, BBX21 positively regulates *HY5*/HY5 at both transcriptional and post-translational levels, directly inducing *HY5* gene expression and forming heterodimers resulting in enhanced HY5 protein activity (Xu et al., 2016; Xu et al., 2018; Bursch et al., 2020; Xu, 2020; Zhao et al., 2020).

While red light signaling enhances nodulation in a *PHYB* dependent manner in the model legume *Lotus japonicus* (Lotus) (Suzuki et al., 2011), blue light exhibits contrary effects on soybean nodulation (Wang et al., 2021; Ji et al., 2022). Soybean *TGACG-MOTIF BINDING FACTOR 3/4* (*GmSTF3/4*), putative orthologues of Arabidopsis *HY5 HOMOLOG* (*AtHYH*), promote nodulation systemically (Wang et al., 2021). In contrast, *GmSTF1/2* and *LjASTRAY*, putatively orthologous to Arabidopsis *HY5,* negatively regulate nodulation in soybean and Lotus, respectively (Nishimura et al., 2002a; Ji et al., 2022). Interestingly, putative legume orthologues of Arabidopsis *HY5* are distinct from the latter in that they possess an additional N-terminal Ring-finger domain and acidic region, both absent in *AtHY5* (Nishimura et al., 2002a) and found only but consistently in the papilionoid lineage (Ji et al., 2022). Consistent with the larger protein size, *Gm*STF1/2 were reported to lack systemic mobility (Chen et al., 2016; Ji et al., 2022).

Beyond the induction of new organs, light also controls the functionality and turnover of established tissue (Sakuraba, 2021). Prolonged periods of darkness induce leaf senescence in several species in a *PHY* and *CRY* dependent manner (Sakuraba et al., 2014; Sakuraba, 2021; Kozuka et al., 2023). Darkness-induced leaf senescence was shown to require *PHYTOCHROME*-*INTERACTING FACTORs* (*PIFs*), which directly induce the expression of *ORESARA1* (*ORE1*) (Sakuraba et al., 2014). In Arabidopsis, the NAC-type transcription factors *ORE1*, *ORE1 SISTER1* (*ORS1*), *NAC-LIKE, ACTIVATED BY AP3/PI* (*NAP*) and *NAC046* have been described as key regulators of leaf senescence (Guo and Gan, 2006; Balazadeh et al., 2011; Qiu et al., 2015; Oda-Yamamizo et al., 2016), whose transcripts accumulate in darkness (Sakuraba et al., 2014; Kozuka et al., 2023). Blue light, on the contrary, counteracted senescence in Arabidopsis *(Kozuka et al., 2023)*. Interestingly, this was partially dependent on *AtHY5* (Kozuka et al., 2023). This apparent role of *HY5* in maintaining organ vitality under light by repressing senescence-related signaling inspired us to investigate a possible role of *HY5* in light-dependent regulation of symbiotic organ vitality. Our study reveals that in Lotus interacting with the rhizobial symbiont *Mesorhizobium loti*, *HY5* along with its putative regulator *BBX21* is required and sufficient to maintain nodule functionality by preventing *NAC094*-dependent light-induced nodule senescence. It thereby links the control of nodule number and –lifetime to light perception as a readout of photoassimilate availability.

## Main

### *LjHY5* and *LjBBX21* prevent premature nodule senescence

We first set out to test whether HY5 mediated light signaling plays a role in symbiosis regulation. Mutant plants carrying an exonic *LORE1* retrotransposon insertion in *Ljhy5-2*, the putative Lotus ortholog of *AtHY5,* exhibited longer primary roots (Fig. 1a) but smaller nodules compared to wild type plants (Fig. 1b-d). Notably, most mature *hy5-2* nodules showed dark discolorations of varying extent (Fig. 1c,d,f). Reddish *hy5-2* nodules lacking such discolorations were in tendency paler than wild type nodules (Fig. 1b,d). To further investigate the localization and appearance of discoloration in *hy5-2* nodules, we sectioned fresh nodule tissues showing varying degrees of blackening (Fig. 1g-j). Sections of both wild-type and *hy5-2* nodules without apparent discoloration showed a typical red interior (Fig. 1g,h). In contrast, in fully or partially discolored *hy5-2* nodules, the bacterially infected nodule cortex included dark brown to black tissue patches (Fig. 1i-j). These appeared to originate in the outer nodule cortex (Fig. 1i) and, in more advanced stages, encompassed the entire extent of the infected nodule center (Fig. 1j).

**Figure 1.**
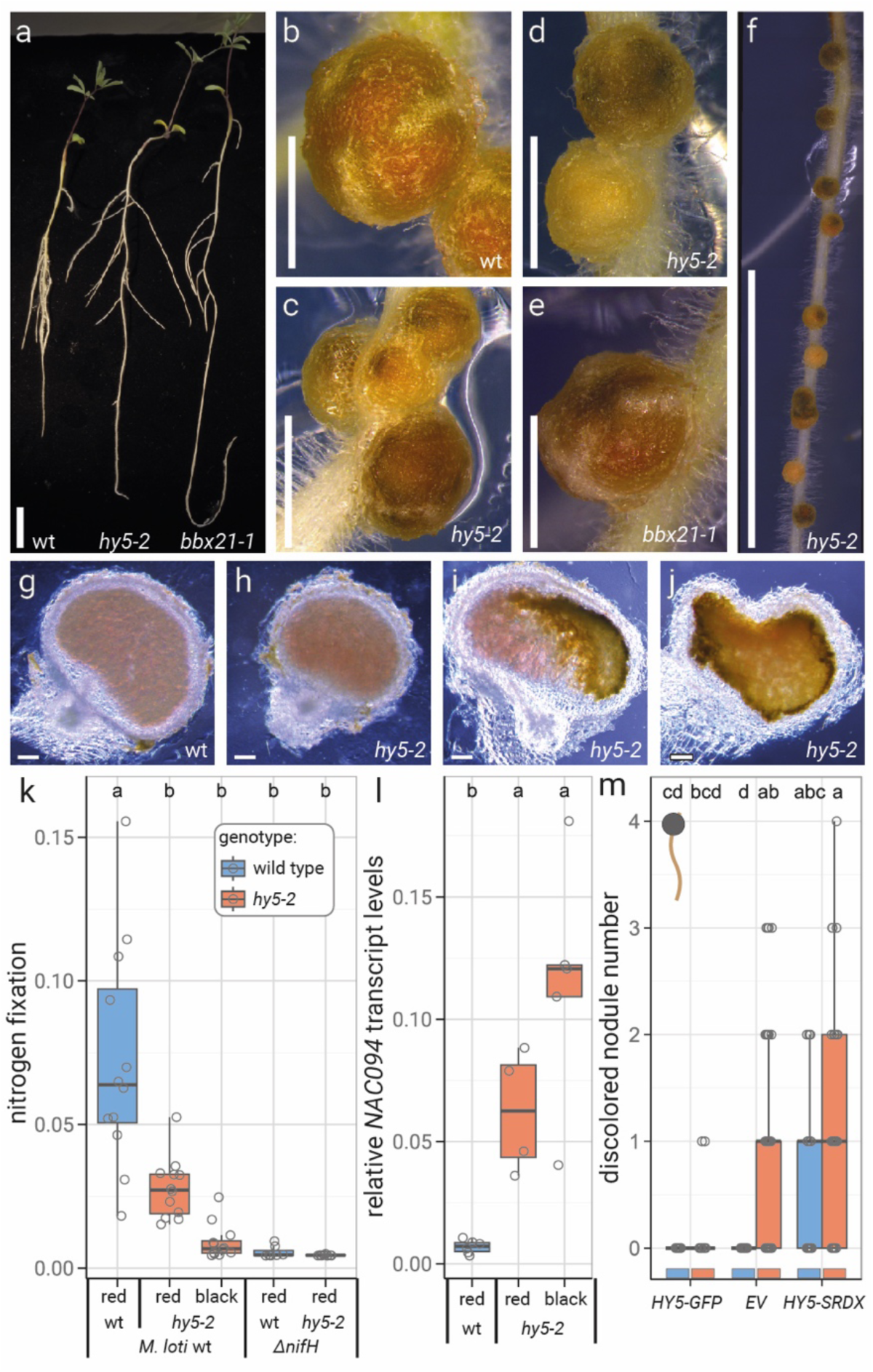
*HY5* and *BBX21* inhibit *NAC094* expression and early nodule senescence. **a**, Phenotype of *L. japonicus* wild type ecotype Gifu (wt), *hy5-2* and *bbx21-1* plants. **b**-**e**, Close-up of wt (**b**), *hy5-2* (**c**-**d**) and *bbx21-1* (**e**) nodules. **f**, Close-up of a *hy5-2* nodule section showing discolored and wt-like nodules. **g**-**j**, Hand sections of wt (**g**) and *hy5-2* nodules displaying different degrees of discoloration (**h**-**j**). Scale bars equal 1 cm (**a**, **f**), 1 mm (**b-e**) or 0.1 mm (**g**-**j**). **k**, Nitrogen fixation in wt and *hy5-2* nodules as determined *via* acetylene reduction. **l**, Relative *NAC094* transcript levels in wt and *hy5-2* nodules. **k**, **l**, Pools of five nodules were harvested for analysis, where red = mature nodules (no apparent discoloration) and black = senescent nodules with varying degrees of dark discolorations. **k**, *ΔnifH* infected control nodules were all pale in color. **m**, Discolored nodule numbers of transiently transformed wt and *hy5-2* roots. **m**, Plants were transformed with either *pUBI1:HY5-GFP* (HY5-GFP), *pUBI1:HY5-SRDX* (HY5-SRDX) or pUBI1:GFP (EV) using *Agrobacterium rhizogenes* and analyzed at 28 days post inoculation with *M. loti* (dpi). **l**, Transcript levels measured *via* qPCR, relative to transcript levels of two housekeeping genes *ATPs* and *PP2a*. **a**-**l**, Plants were evaluated at 21 dpi. Comparisons used Analysis of Variance (ANOVA) and *post-hoc* Tukey testing (p≤0.05), with distinct letters indicating significant differences p≤0.05. Boxplot central line shows median value, box limits indicate the 25^th^ and 75^th^ percentiles. Whiskers extend 1.5 times the interquartile range or to the last datapoint. Individual data points are represented by dots.

Acetylene reduction assays revealed reduced nitrogenase activity in both reddish and partially or fully discolored *hy5-2* compared to mature red wild type nodules (Fig. 1k). Nitrogen fixation in discolored *hy5-2* nodules by wild type *M. loti* resembled that in nodules infected with an *M. loti* Δ*nifH* strain (Ooki et al., 2005) unable to induce nitrogen-fixing nodules (Ooki et al., 2005; Fotelli et al., 2011) (Fig. 1k). These results demonstrate that nodules formed on *Ljhy5-2* mutant plants are significantly impaired in nitrogen fixation, with those nodules exhibiting dark discoloration showing nitrogenase activity levels near the detection limit (Fig. 1k).

Localized dark discoloration was previously proposed to correlate with activated defense signaling in both Lotus (Goto et al., 2025) and *Medicago truncatula* (Berrabah et al., 2024) nodules. We thus compared transcript levels of the defense associated genes *PR8* (Berrabah et al., 2023) and *PR10* (Nakatsukasa-Akune et al., 2005; Berrabah et al., 2023) in wild type and *hy5-2* nodules. Independent of genotype and nodule phenotypic appearance, we observed similar *PR8* and *PR10* transcript abundances (Supp. Fig. 1a,b), suggesting that the defense status of wild type and *hy5-2* nodule tissue is not significantly different. The observed discolorations and reduced nitrogenase activity of *hy5-2* nodules are thus unlikely to primarily reflect an altered immune status.

Inspired by the known role of Arabidopsis *HY5* in counteracting leaf senescence, we then investigated whether the impaired nitrogen fixation efficiency and discoloration of nodules formed in the absence of *HY5* are linked to senescence processes. Along this line, we quantified transcript levels of the senescence associated transcription factor *NAC094* in *hy5-2* mutant nodules. qPCR-based expression analysis revealed significantly increased *NAC094* transcript levels in *hy5-2* compared to wild type nodules independent of coloration (Fig. 1l; Supp. Fig. 1c).

To verify that loss of *HY5* functionality in *hy5-2* mutants is causative of senescent nodule formation, we overexpressed the Lotus *HY5* gene coupled to a visual marker in *hy5-2* roots, resulting in an almost complete absence of early senescent nodules (Fig. 1m). Notably, when we fused *HY5* to a transcriptional repressor motif, *EAR repressor motif* of the Arabidopsis *SUPERMAN* gene (SRDX) (Hiratsu et al., 2003), this triggered the formation of senescent nodules in wild type as well as *hy5-2* mutants (Fig. 1m), mimicking the *hy5-2* loss-of-function phenotype (Fig. 1c,d,f). This *SRDX* motif was previously shown in Arabidopsis to mediate repression of *HY5* target genes that are activated by wild type *HY5* (Burko et al., 2020). In summary, these data suggest that in the absence of a functional *HY5* gene, nodules enter premature senescence, reflected by enhanced *NAC094* activity and darkening of nodule tissue, correlated with diminished nitrogen fixation. In Lotus, *HY5* thus seems to be essential in maintaining nodule function and vitality, counteracting the onset of senescence in healthy plants. Complementarily, expression of the dominant negative *HY5-SRDX* fusion is sufficient to induce nodule senescence.

In Arabidopsis, *HY5*/HY5 gene/protein activity underlies joint regulation by several *BBX* gene products (Bursch et al., 2020; Xiao et al., 2021). Among these, *AtBBX21* plays a key role. It activates *AtHY5* transcription (Xu et al., 2016; Xu et al., 2018) and acts as a cofactor of the *At*HY5 protein, mediating transcriptional control of its target genes (Datta et al., 2007; Bursch et al., 2020; Xiao et al., 2021). An investigation of *BBX* transcription factors in Lotus identified 23 *BBX* gene homologs in the *Lotus japonicus* genome (v1.3) (Supp. Fig. 2a) (Kamal et al., 2020). Further comparative analysis of gene/protein sequences revealed LotjaGi5g1v0010400 (*Lj*BBX21) as putative orthologue of *At*BBX21 (Supp. Fig. 2a). *At*BBX21 and *Lj*BBX21 amino acid sequence alignments reveal a high degree of conservation in predicted functional protein domains. These include BBOX1, BBOX2 and COP1 binding domains (Supp. Fig. 2b). Similarly, *At*HY5 and *Lj*HY5 exhibit highly conserved COP1 binding and bZIP domains (Supp. Fig. 2c). These data suggest that at the protein level, HY5 and BBX21 may display similar regulatory activity patterns in Arabidopsis and Lotus. To test this, we isolated a mutant line carrying an exonic LORE1 retrotransposon insertion in the putative *LjBBX21* gene locus (*bbx21-1*) (Malolepszy et al., 2016). Root and nodule development and appearance in *bbx21-1* resembled *hy5-2* plants (Fig. 1 a,c-e). This is in line with a partial dependency of *HY5/*HY5 gene/protein activity on BBX21 in Lotus. Like *hy5-2*, *bbx21-1* mutants showed enhanced primary root growth compared to wild type (Fig. 1a), and *bbx21-1* nodules frequently showed dark discoloration (Fig. 1e). *NAC094* transcript levels in *bbx21-1* nodules were intermediate, ranging between levels observed in wild type and *hy5-2* (Supp. Fig. 1c). These data are in line with a scenario where *BBX21* regulates the function of *HY5* in maintenance of nodule function and viability.

### *BBX21* and *HY5* act in concert to protect nodules from light induced senescence

To investigate whether Lotus *BBX21* and *HY5* act in the same genetic pathway, we generated double mutant lines using *hy5-2* and *bbx21-1* genetic backgrounds. Indeed, *hy5-2 bbx21-1* double mutants developed nodule numbers similar to *hy5-2* single mutants (Fig. 2a-b). Consistently, the senescence status of mature nodules as judged by *NAC094* transcript levels (Fig. 2c) and nitrogen fixation rates (Fig. 2d) resembled that of *hy5-2* single mutants. This suggests that *HY5* acts epistatically to *BBX21* in Lotus. These results are consistent with reports that in Arabidopsis, BBX21 regulates *HY5*/HY5 activity (Xu et al., 2016; Xu et al., 2018; Zhao et al., 2020). Analysis of transcript accumulation levels revealed that *HY5* transcripts are wild type-like in both roots and nodules of Lotus *bbx21-1* mutant plants (Supp. Fig. 3a,b). Similarly, *BBX21* transcript levels were wild type-like in *hy5-2* mutants (Supp. Fig. 3c,d). These data suggest that that HY5 and BBX21 work in the same genetic pathway, where regulatory activity likely occurs at the protein level rather than *via* transcriptional control.

**Figure 2.**
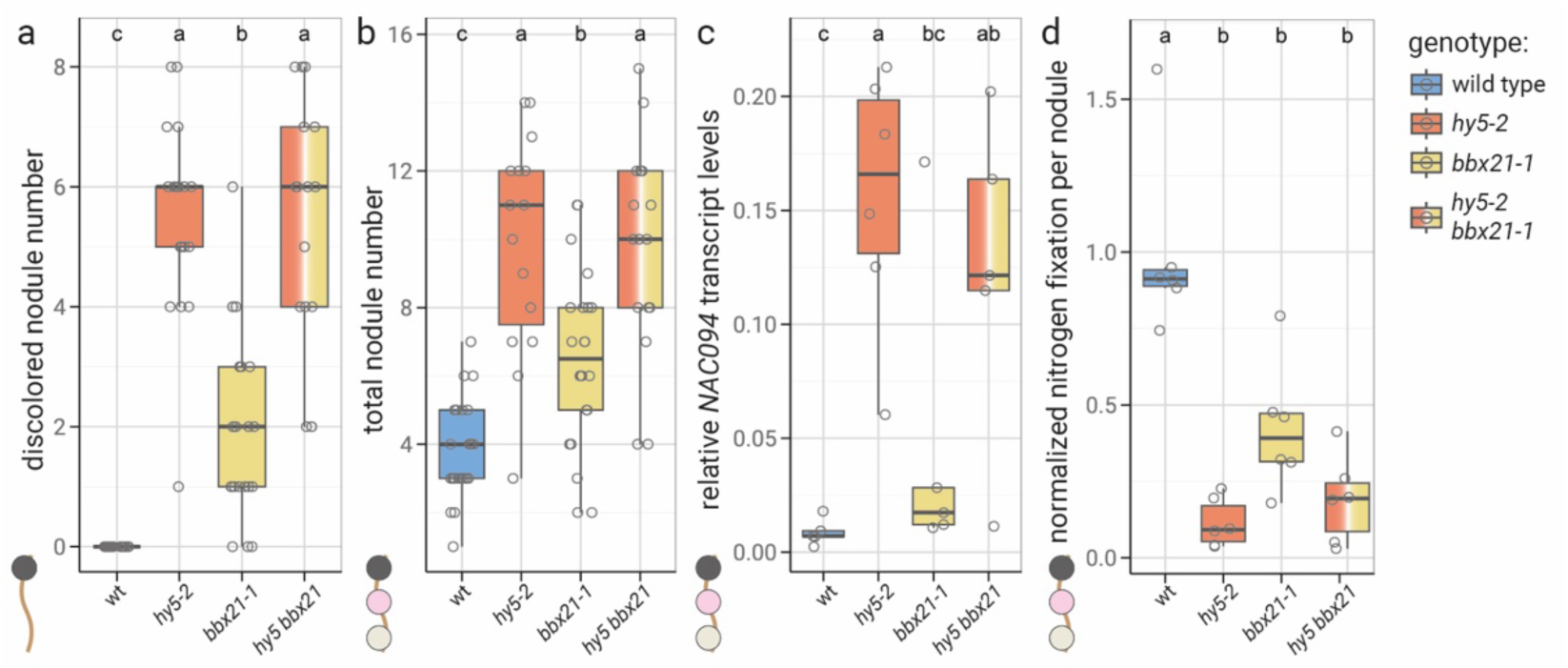
*HY5* and *BBX21* act in concert to guard nodules from light induced senescence. **a**-**d**, Discolored nodule numbers (**a**), total nodule numbers (**b**), relative *NAC094* levels (**c**) and nitrogen fixation per nodule (**d**) of *L. japonicus* Gifu wild-type (wt), *hy5-2*, *bbx21-1* and *hy5-2 bbx21-1* double mutant plants. Plants were grown under white light (200 µE) and evaluated three weeks post inoculation. Transcript accumulation levels are relative to two housekeeping genes (*ATPs* and *PP2A*). Comparisons used Analysis of Variance (ANOVA) and *post-hoc* Tukey testing (p≤0.05), with distinct letters indicating significant differences p≤0.05. Boxplot central line shows median value, box limits indicate the 25^th^ and 75^th^ percentiles. Whiskers extend 1.5 times the interquartile range or to the last datapoint. Individual data points are represented by dots.

In Arabidopsis, *At*BBX20/21/22 function as rate limiting cofactors of *At*HY5 at the protein level (Bursch et al., 2020). Our combined data are consistent with a scenario where Lotus *Lj*BBX21 similarly acts as cofactor of *Lj*HY5, mediating *Lj*HY5 dependent nodule viability maintenance. The consistently intermediate phenotype of *Ljbbx21-1* averaging between wild type and *Ljhy5-2* plants suggests a possible involvement of additional *Lj*BBX proteins in regulating *Lj*HY5 activity.

### Root *HY5* and *BBX21* regulate nodule functionality and number

A Lotus mutant line retaining the second *HY5* intron, *Ljsym77*/*astray/Ljhy5-1*, has been reported earlier to show agravitropic lateral roots and a hypernodulation phenotype (Nishimura et al., 2002b; Nishimura et al., 2002a). Consistently, both *hy5-2* and *bbx21-1* plants develop an overabundance of nodules compared to wild type (Fig. 3a).

**Figure 3.**
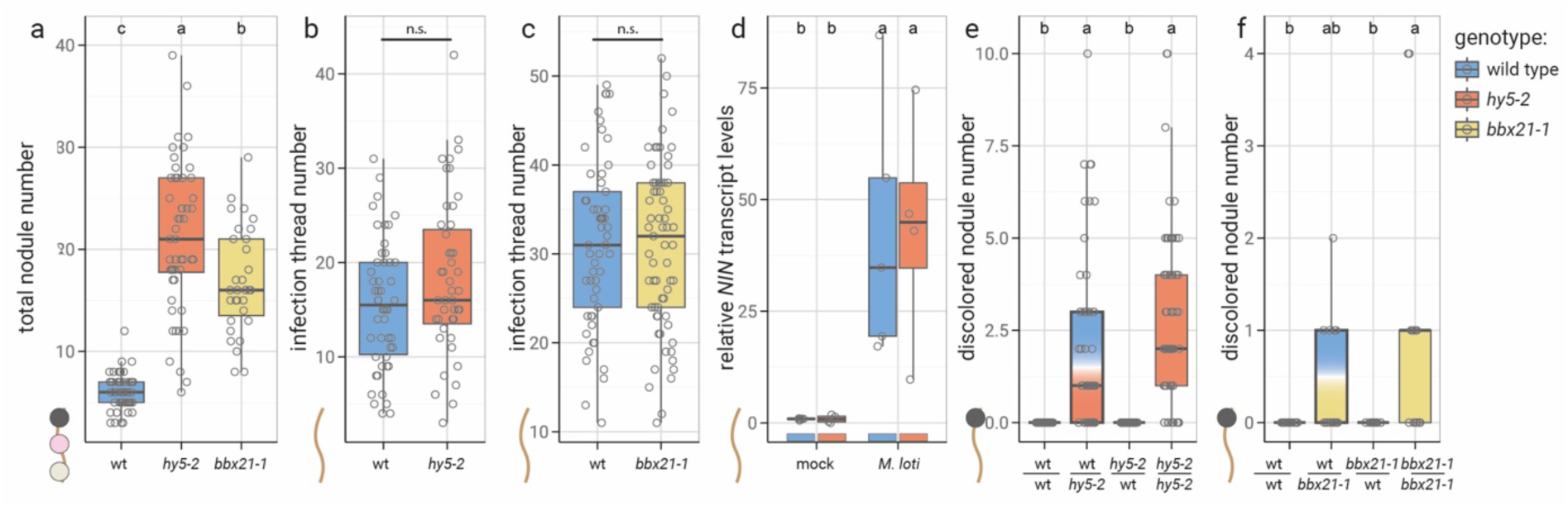
Nodule function and number, but not infection, depend on root *HY5* and *BBX21*. **a**, Total nodule numbers of Gifu wild type (wt), *hy5-2* and *bbx21-1* plants. **b**, **c,** Number of infection threads *hy5-2* (**b**) and *bbx21-1* compared to wt plants (**c**) inoculated with *M. loti*. **d,** Relative *NIN* levels in *L. japonicus* roots three days post infection. Transcript levels were determined *via* qPCR, relative to two housekeeping genes. **a-e,** Plants evaluated after 1 week (**b**, **c**), two weeks (**d**) or three weeks (**a**) of growth in sterile culture. **e**, **f,** Numbers of discolored nodules on grafted plants three weeks post inoculation with *M. loti*. Comparisons used Student’s t-test (**b**, **c**) or Analysis of Variance (ANOVA) and *post-hoc* Tukey testing (p≤0.05) (**a**, **d**-**f**), with distinct letters indicating significant differences p≤0.05. Boxplot central line shows median value, box limits indicate the 25^th^ and 75^th^ percentiles. Whiskers extend 1.5 times the interquartile range or to the last datapoint. Individual data points are represented by dots.

Apart from hypernodulation, soybean loss-of-function mutants of the putative *HY5* orthologues *STF1/2* show enhanced infection thread numbers (Ji et al., 2022). In line with this, soybean *STF1/2* genes appear to restrict nodulation symbiosis via repression of *NODULE INCEPTION* (*NIN)* (Ji et al., 2022), a gene required both for infection and organogenesis during RNS (Schauser et al., 1999; Soyano et al., 2013). Interestingly, nodule blackening was not reported in *Ljhy5-1* (Nishimura et al., 2002b; Nishimura et al., 2002a), nor in *stf1 stf2* mutants in soybean (Ji et al., 2022).

In contrast to soybean, Lotus *hy5-2* and *bbx21-1* mutants both showed wild-type like infection phenotypes (Fig. 3b,c). Consistently, *NIN* transcript accumulation at 3 days post inoculation (dpi) with *M. loti* was similar in *hy5-2* and wild-type plants (Fig. 3d) These combined data suggest that in Lotus, epidermal infection and early symbiotic development are *HY5*-independent.

To investigate whether *HY5* and *BBX21* dependent regulation of nodule establishment and senescence in Lotus is local or systemic, we conducted grafting experiments. Discolored nodules were exclusively found on plants possessing *hy5-2* rootstocks (Fig. 3e). In contrast, nodules on plants with *hy5-2* shoots on wild-type root systems showed no signs of blackening (Fig. 3e). A similar trend was observed in grafts of wild-type and *bbx21-1* plants (Fig. 3f). Consistently, nodules on grafts with *hy5-2* or *bbx21-1* rootstocks showed significantly reduced nitrogen fixation rates compared to grafts with wild type root stocks (Supp. Fig. 4a). At the same time, hypernodulation was linked to *hy5-2* or *bbx21-1* rootstocks, whereas plants with respective mutant shoots developed wild type-like nodule numbers if grafted on wild type roots (Supp. Fig. 4b-e). These combined data suggest that *HY5* and *BBX21* act in roots both to inhibit nodule senescence and to restrict nodule organogenesis events. Double mutant analyses suggest that *HY5* acts largely epistatically in the observed functional context. We thus focused our further investigations on *hy5* mutant plants.

### Illumination triggers early nodule senescence and nodule proliferation in the absence of *HY5*

As *HY5* has been associated with photomorphogenesis in Arabidopsis (Osterlund et al., 2000; Xiao et al., 2021), we hypothesized that Lotus *HY5* may regulate nodule senescence in dependence of light intensity, an indicator of photoassimilate availability. To investigate this, we grew wild-type and*hy5-2* plants at different white light intensities, with their roots shielded from direct light exposure. Independent of light intensity, wild-type plants did not exhibit nodules with apparent signs of senescence after three weeks of co-cultivation (Fig. 4a). In contrast, *hy5-2* mutants developed discolored, putatively senescent nodules in a light dosage dependent manner (Fig. 4a). Interestingly, at low light intensities, discolored nodules were absent also on *hy5-2* plants (Fig. 4a), identifying the formation of discolored nodules in this genetic background as conditional. Light conditions may thus explain the fact that discoloration has not previously been described in legume *hy5/stf* mutants (Nishimura et al., 2002b; Nishimura et al., 2002a; Ji et al., 2022).

**Figure 4.**
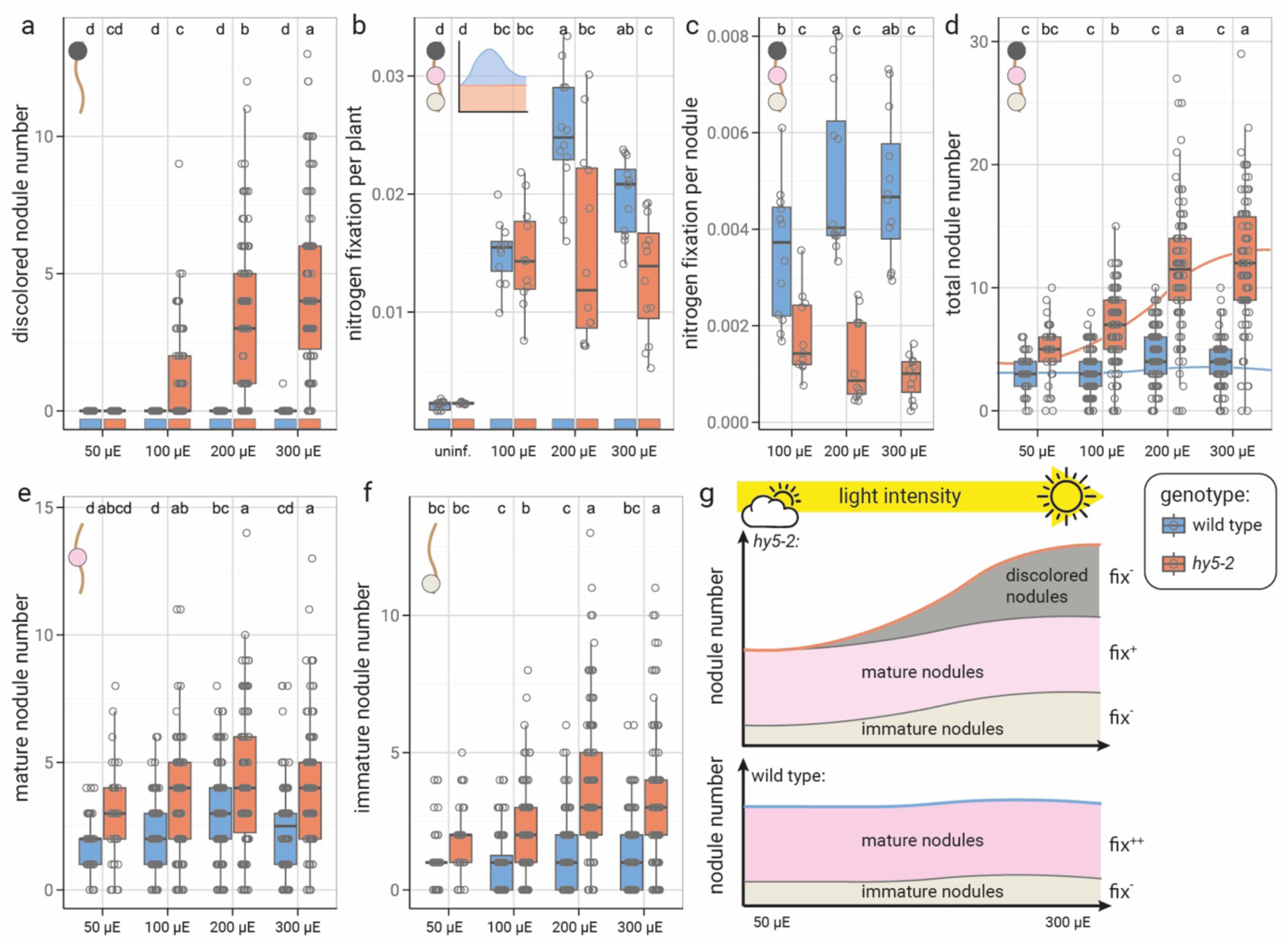
*HY5* inhibits light dependent nodule senescence. **a**-**f,** Nodule numbers and nitrogen fixation of *L. japonicus* Gifu wild-type (wt) and *hy5-2* plants. **a**, **d**-**f,** Discolored nodule number (**a**), total nodule number (**d**), mature, non-senescent nodule number (**e**) and immature nodule number (**f**) of the same plants. **b**, **c** Nitrogen fixation per plant (**b**) and nitrogen fixation per nodule (**c**) of a subset of plants evaluated in **a**, **d**-**f.** Plants evaluated three weeks post infection. Plants were grown at indicated light intensities of white light, roots were shielded from light exposure. Comparisons used Analysis of Variance (ANOVA) and *post-hoc* Tukey testing (p≤0.05) with distinct letters indicating significant differences p≤0.05. Boxplot central line shows median value, box limits indicate the 25^th^ and 75^th^ percentiles. Whiskers extend 1.5 times the interquartile range or to the last datapoint. Individual data points are represented by dots. **g,** Model of the light dependency of nodule formation, maturation and senescence in wild-type and *hy5-2* plants.

To evaluate how light intensity and light-triggered senescence affect nodule function, we compared nitrogen fixation efficiency between light conditions. Nitrogen fixation in wild-type plants peaked at intermediate light intensity (200 µE; Fig. 4b-c). Consistently, while overall nodule numbers remained similar (Fig. 4d-g), wild type plants supported more mature nodules at 200 µE than at other light intensities (Fig. 4e). In addition, fixation capacity of mature nodules positively correlated with light availability (Supp. Fig. 5).

At low light, *hy5-2* mutants fixed nitrogen at similar levels as wild-type plants (Fig. 4b). However, in the absence of *HY5*, fixation capacity of plants and nodules did not rise with enhanced light availability (Fig. 4b-c; Supp. Fig. 5). In contrast, relative to wild type, fixation capacity of *hy5-2* mutant nodules was increasingly compromised with rising light intensity (Supp. Fig. 5a).

The combined data suggest that in a wild type-situation, nitrogen fixation is optimal at intermediate ambient light intensities around 200 µE, with both lower and higher intensities yielding reduced fixation levels per plant. Our data suggest that this primarily reflects *HY5*-dependent light responsiveness of fixation capacity at the nodule level (Fig. 4 b,c; Supp. Fig. 5). Interestingly, the peak in mature nodule numbers found in wild-type plants at 200 µE was not observed in *hy5-2* plants, despite a significant increase in both discolored and immature as well as overall nodule abundance with rising light intensity in the mutant (Fig. 4a, d-g). The light-dependent accumulation of discolored (Fig. 4a,g) as well as immature (Fig. 4f-g) nodules in *hy5-2* mutants is consistent with a scenario where in the absence of *HY5*, mature nodules shift towards a senescent state in a light intensity-dependent manner, paralleled by a release of AON and thereby the development of new nodule primordia. Notably, these combined data suggest that a mechanism balancing nodule numbers with the plants’ overall nitrogen status is in place also in the absence of *HY5*. Evidently, this mechanism is epistatic to – and possibly causative of – the *hy5-2* hypernodulation phenotype.

Consistent with this hypothesis, other Lotus mutants developing dysfunctional nodules impaired in nitrogen fixation similarly show enhanced nodule proliferation. For example, mutants defective in the *SYMBIOSIS SULFATE TRANSPORTER 1* (*SST1*) gene (Krusell et al., 2005) or the putative iron transporter gene *STATIONARY ENDOSYMBIONT NODULE 1* (*SEN1*) (Suganuma et al., 2003; Hakoyama et al., 2012; Walton et al., 2020) show early nodule senescence paralleled by the continuous formation of new nodule organs. As a result, *sst1* and *sen1* mutants, like *hy5-2*, are hypernodulating (Suganuma et al., 2003; Krusell et al., 2005).

### Red and blue light affect nodule lifespan and function both systemically and locally

To decipher which light fraction may be causative of nodule senescence and hypernodulation in *hy5-2* mutants, we selectively exposed plants to blue light, red light or combined red and far-red light (Supp. Fig 6a-f). Simulating natural growth conditions, we initially irradiated above-ground plant parts only, keeping roots shaded from light. Wild-type plants formed a similar number of nodules under all tested light regimes (Fig. 5a). None of these coincided with the formation of discolored or dysfunctional nodules (Fig. 5b-c). Relative to wild type, *hy5-2* mutants were consistently hypernodulated (Fig. 5a). Blue and red light triggered a similar nodule number increase compared to wild type plants (Fig. 5a), coinciding with the occurrence of senescent nodules (Fig. 5b). Blue light was most effective in inducing nodule senescence (Fig. 5b) and also proved the strongest inhibitor of nitrogen fixation (Fig. 5c). Notably, far-red light partially rescued red light-induced hypernodulation and senescence induction in *hy5-2* (Fig. 5a,b,d). This is consistent with a role of PHYTOCHROME B (PHYB) in red light mediated symbiosis control. PHYB is activated by red light but inactivated by far-red light (Suzuki et al., 2011; Legris et al., 2019). Numbers of mature, reddish nodules did not differ significantly between those light conditions for neither wild-type, nor *hy5-2* plants (Supp. Fig. 7a). As light was applied to shoots only, the observed effects on root symbiosis must reflect systemic processes. Based on grafting results (Fig. 3e-f), *HY5* and *BBX21* are predicted root mediators translating shoot light signaling into root symbiotic responses and preventing nodule senescence. Thus, both blue and red light perception by shoots seems sufficient to trigger root nodule abortion in the absence of functional *HY5* or *BBX21* loci. This reveals the existence of a shoot-derived systemic signal that activates root *HY5*/*BBX21* mediated maintenance of nodule vitality upon red or blue light perception in the shoot. Possible candidates for blue light dependent systemic signals could be *HYH* or *FT* homologues, as these genes have been shown to be involved in systemic blue light signaling downstream of *CRY1* in soybean (Wang et al., 2021).

**Figure 5.**
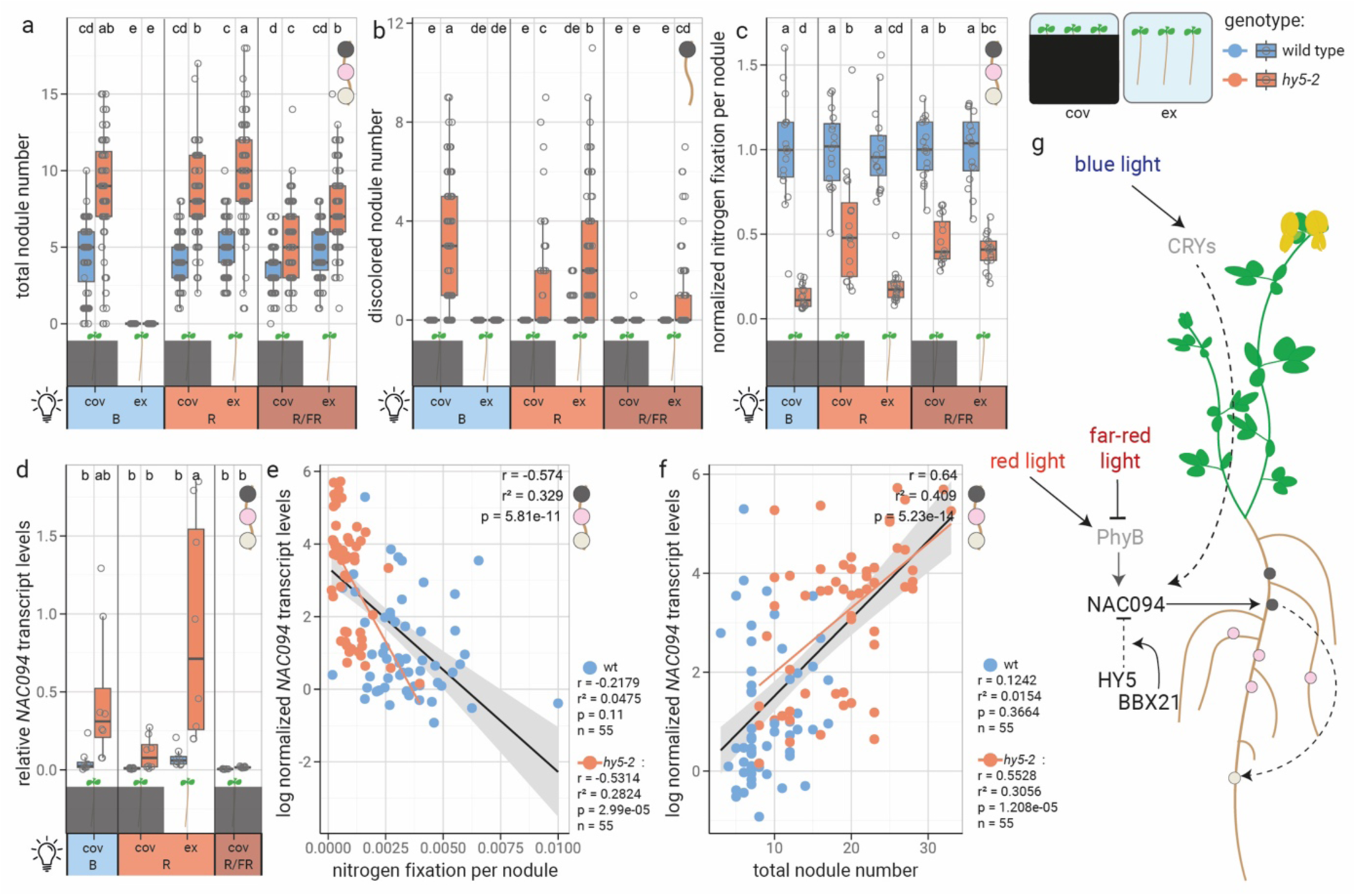
Light controls *NAC094*-induced nodule senescence in a wavelength-dependent manner. **a**, **b,** Number of discolored nodules (**a**) and total nodules (**b**) of the same plants. **c,** Normalized nitrogen fixation per nodule of a subset of plants evaluated in (**a**, **b**). Data were normalized to the average fixation per nodule of wild-type plants. Plants were analyzed at 21 days post inoculation with *M. loti*. **d,** Relative *NAC094* expression in nodules harvested of a subset of plants used in (**c**). Expression was determined relative to two *ATPs* and *PP2A*. Plants evaluated three weeks (**a**-**d**) post infection with *M. loti*. Plants were grown at indicated light regimes at 200 µE light intensity. R = red light (λ=660 nm); R/FR = red light (λ=660 nm) with far-red supplementation (λ=730 nm); B = blue light (λ=450 nm); cov = roots covered from light exposure; ex = roots exposed to light. **e**, **f,** Correlation plots. Analyses include data shown in **Fig. 2d**, **Supp. Fig. 5b**; **Fig. 5d** and **Supp. Fig. 1c**. Pearson correlation analysis was conducted for each dataset as a whole (datapoints of all genotypes were pooled) and for genotype-specific subgroups. Respective results parameters of the genotype specific analyses are displayed next to the graphs. Where a significant correlation (p<0.05) was observed, trendlines are displayed. For genotype specific data, the color code is as indicated to the right of the graphs. Where datapoints of all genotypes were pooled, trendlines are in black, with grey areas indicating 95% variance limits. log, natural logarithm. **a**-**d,** Comparisons used Analysis of Variance (ANOVA) and *post-hoc* Tukey testing (p≤0.05) with distinct letters indicating significant differences p≤0.05. All data presented in the respective panels was included in the analyses, vertical separators are visual aids only. Boxplot central line shows median value, box limits indicate the 25^th^ and 75^th^ percentiles. Whiskers extend 1.5 times the interquartile range or to the last datapoint. Individual data points are represented by dots.

We then released root shading to determine whether beyond its systemic role, light might directly and locally impact root symbiotic performance. Notably, direct blue light access to roots inhibited nodulation entirely (Fig. 5a) (Shimomura et al., 2016). It further led to impaired growth as well as overall plant performance (Supp. Fig. 6f), identifying blue light as a potent inhibitor of plant development if (co-)perceived by roots. Similarly, *hy5-2* mutant plants were asymbiotic when roots were exposed to blue light. Selective red-light irradiation, in contrast, did not affect nodule formation in wild-type plants, but led to enhanced nodule numbers in *hy5-2* mutants if applied to roots (Fig. 5a-b). Consistently, average nitrogen fixation rates of *hy5-2* plants plummeted upon root exposure to red light (Fig. 5c; Supp. Fig. 7b,c), along with *NAC094* transcript accumulation compared to plants where only shoots were illuminated (Fig. 5d, Supp. Fig. 7d). As previously observed for shoot mediated, systemic effects, root illumination with far red light in addition to red light counteracted red light effects on roots (Fig. 5a-c).

In summary, the formation of discolored nodules, high nodular *NAC094* levels and low nitrogen fixation were consistent characteristics of *hy5-2* mutant compared to wild type plants (Fig. 1a-m; Fig. 2a-d; Fig. 4a-c; Fig. 5a-d). In line with this, correlation analysis (Fig. 5e,f; Supp. Fig. 8a-f) revealed a significant correlation between *NAC094* levels and senescent nodule number for *hy5-2*, but not wild type plants (Supp. Fig. 8a). Also, *NAC094* expression levels were negatively correlated with nitrogen fixation activity in nodules in *hy5-2* (Fig. 5e). Remarkably, *hy5-2* total nodule numbers correlated with *NAC094* transcript levels and the occurrence of discolored nodules (Fig. 5f; Supp. Fig. 8b). Consistently, overall nitrogen fixation correlated with nodule numbers in wild type, but not in *hy5-2* mutant plants (Supp. Fig. 8c). We observed similar interrelations in *bbx21-1* plants (Supp. Fig. 8a-e). In line with previous observations, these correlations were consistently weaker compared to *hy5-2* plants, suggesting that other factors may co-regulate HY5 in a partially redundant fashion. These data support the hypothesis that hypernodulation in *hy5-2* and *bbx21-1* mutants may reflect a release of AON due to insufficient nitrogen fixation as a result of *NAC094* induced senescence. In line with this hypothesis, *NAC094* overexpression lines were previously reported to hypernodulate, and fail to produce functional nodules (Wang et al., 2023). We propose *HY5* dependent repression of *NAC094* to be indirect, as overexpression of a dominant negative *HY5-SRDX* construct was sufficient to trigger nodule senescence in wild-type plants (Fig. 1m).

Collectively, our findings reveal roles of *HY5* and *BBX21* in safeguarding nodules from early senescence and ensuring a robust and efficient nitrogen fixation at various light conditions (Fig. 5g). This establishes the control of nodule function and lifespan as a critical component in symbiosis regulation, and in balancing symbiosis in the light of dynamic abiotic growth conditions.

### Perspective

In the asymbiotic Arabidopsis, HY5 is a master regulator of light signaling, and integrates light availability and photomorphogenesis with senescence processes and key physiological domains such as nutrient homeostasis (Xiao et al., 2021; Mankotia et al., 2024).

These reported activities are consistent with a role of legume HY5 in protecting nodule organs from light-induced senescence and integrating light availability with nodule functionality. Beyond a systemic role in carbon-nitrogen balance and nitrate uptake control (Chen et al., 2016), root HY5 promotes and regulates sulfate, iron and phosphate acquisition and processing (Xiao et al., 2021; Mankotia et al., 2024). Nodule organs represent a strong sink for these nutrients, as iron and sulfate are essential components of the active nitrogenase complex (Zanello, 2019), and their availability is a prerequisite for nodule functionality. A failure of bacteroid-containing symbiosomes to internalize sulfate (*sst1*) and iron (*sen1*) from the host plant cell renders nodules unable to fix nitrogen (Suganuma et al., 2003; Krusell et al., 2005). The observation that Lotus *hy5-2* mutants are severely impaired in nitrogen fixation may thus be linked to limitations in macronutrient availability. As in *hy5-2* mutants, a failure to fix nitrogen in *sen1* and *sst1* plants coincides with early nodule senescence and a hypernodulation phenotype (Suganuma et al., 2003; Krusell et al., 2005).

In Arabidopsis, *HY5* is further involved in the protection of both roots (Zhu et al., 2025) and above-ground tissues (Wang et al., 2018) from oxidative stress. It transcriptionally represses *MYB30*, a central mediator of oxidative stress responses (Burko et al., 2020; Zhu et al., 2025). *Athy5* mutants not only displayed hypersensitivity to hydrogen peroxide (H_2_O_2_) but also accumulated H_2_O_2_ in root apical meristems (Zhu et al., 2025). A possible disbalance of redox homeostasis and raised H_2_O_2_ levels in Lotus *hy5-2* mutant nodules may influence the redox state of leghemoglobin associated iron atoms. In the presence of trace amounts of peroxides, oxygenated ferrous leghemoglobin (containing Fe^2+^) readily auto-oxidizes to ferric leghemolobin (Fe^3+^) (Becana and Klucas, 1992). In its ferric state, leghemoglobin is unable to function as O_2_ carrier (Becana and Klucas, 1992) and protect bacterial nitrogenase in infected host cells from oxygen damage, resulting in compromised fixation capacity. Contrasting with the red appearance of active leghemoglobin, the ferric form of the latter further displays a brown color (Virtanen and Laine, 1946; Becana and Klucas, 1992). Superoxide-mediated conversion of functional, ferrous leghemoglobin to a dark brown ferric state may thus explain the observed discoloration as well as compromised functionality of *hy5-2* nodules.

Notably, HY5 depletion has been associated with enhanced resistance to nutrient starvation by fostering autophagy mediated endogenous recycling processes (Yang et al., 2020). It is an attractive hypothesis that HY5 regulation, apart from integrating symbiosis with the overall nutrient and physiological status of the host plant, ensures efficient re-use of nutrients delivered to symbiotic nodule organs along with mediating their controlled decline.

From an evolutionary perspective, the recruitment of the *HY5* master regulator displaying an extreme level of regulatory interconnectivity and a significant fitness relevance is a fascinating example of functional diversification. Even minor shifts in HY5 activity are, under the dynamic abiotic conditions of natural growth environments, expected to significantly compromise plant resilience. Evolutionary adoption of *HY5* as a governor of nodule lifetime enabled a dynamic integration of endosymbiotic nutrient exchange in the plants’ global C/N and mineral nutrient balance. Nodulation symbiosis displays a history of genetic neofunctionalizations. The putative recruitment of infection genes from arbuscular mycorrhiza (Kistner and Parniske, 2002), and of organogenesis genes from lateral root formation (Dong et al., 2020; Sexauer et al., 2023; Sexauer and Markmann, 2026) has been well documented. HY5 represents a further stunning example, where functional diversification of existing genes forms a foundation for the acquisition and maintenance of novel features.

## Methods

### Plant and bacterial resources

Plants used in this study were *Lotus japonicus* ecotype Gifu B-129 wild type (Handberg and Stougaard, 1992), and lines carrying exonic LORE1 retrotransposon insertions (Malolepszy et al., 2016) in the genes *HY5* (LotjaGi6g1v0183200) and *BBX21* (LotjaGi5g1v0010400). The mutant lines were referred to as *hy5-2* (line ID 30086963), *bbx21-1* (line ID 30036860) and *bbx21-2* (line ID 30092787), respectively. Further, a *bbx21-1 hy5-2* double mutant line was generated by crossing. Primers used for genotyping of these plants are listed in Supplementary Table 1. Plants were inoculated with *Mesorhizobium loti* strains R7A wild type, MAFF303099 wild type (Kaneko et al., 2000) or MAFF303099 expressing *DsRED* bacteria (Maekawa et al., 2009). Transient transformation of *L. japonicus* plants made use of *Agrobacterium rhizogenes* strain AR1193 (Hansen et al., 1989). Cloning approaches utilized *E. coli* strains DH10b (Grant et al., 1990) or DB3.1 (Bernard et al., 1994).

### Plasmid construction and sequence analysis

Plasmids used for transient transformation of *L. japonicus* are listed in Supplementary Table 2. Plasmid generation followed a modified Golden Gate technology-based approach based on (Weber et al., 2011). *HY5* and *BBX21* coding sequences were amplified from *L. japonicus* (ecotype Gifu) genomic DNA using primers carrying overhangs for subsequent Golden Gate-based cloning. Primer sequences are listed in Supplementary Table 3. *In silico* cloning and sequence analysis was done using CLC Main Workbench 25 (Qiagen). *BBX* gene homologs in the Lotus genome were identified *via* the BLASTN algorithm (Altschul et al., 1990), using Arabidopsis BBX protein sequences as bait. For phylogenetic analysis, sequences were aligned using the ClustalW algorithm (Thompson et al., 1994). The phylogeny was inferred using the Maximum Likelihood method and Jones-Taylor-Thornton model of amino acid substitutions (Jones et al., 1992). The initial tree for the heuristic search was selected by choosing the tree with the superior log-likelihood between a Neighbor-Joining (NJ) tree (Saitou and Nei, 1987) and a Maximum Parsimony (MP) tree. The NJ tree was generated using a matrix of pairwise distances computed using the Jones-Taylor-Thornton model. Evolutionary analyses were conducted in MEGA12 (Kumar et al., 2024).

### Plant growth and infection

For plant growth and phenotyping*, L. japonicus* seeds were surface scarified, sterilized using sodium hypochloride solution containing 1 g/l NaClO, imbibed in ddH_2_O and transferred to sterile growth medium containing 1% (w/v) phyto agar (Duchefa Biochemie) in water. Following stratification at 4 °C (three days), seeds were germinated at 21 °C in constant darkness (three days). For growth on plates, seedlings were transferred to 12 × 12 cm square plastic dishes containing 50 ml ½ –strength modified B&D / 1% (w/v) phyto agar medium without nitrate and inoculated with *M. loti* OD 0.01 via drop inoculation. Modified B&D medium was prepared according to (Broughton and Dilworth, 1971), iron citrate has been switched for Fe-EDTA at 40 µM concentration for ½ strength B&D, pH has been buffered using 4 ml of 0.5M MES pH 6.1 stock per liter of medium. Plants were grown at long day conditions (16 h light, 21 °C / 8 h dark, 17 °C) at variable light conditions indicated in figure legends. Light conditions included a sun like white light spectrum at different light intensities (Polyklima True Daylight plus) or blue light (λ=450 nm), red light (λ=660 nm), or red supplemented with far-red (λ=730 nm) light at 200 µM/s/m^2^. Unless stated differently, plants were grown at 200 µM/s/m^2^ white light.

### Transient transformations

For transient “hairy root” transformation, *L. japonicus* seeds were germinated as described above. Emerging root tips were then pierced into slanted ½ strength Gamborg B5 medium (Gamborg et al., 1968) and transferred to 21°C darkness to induce hypocotyl elongation. In parallel, *A. rhizogenes* carrying desired plasmids for transformation were streaked on LB plates and grown at 28 °C for formation of a dense bacterial film. Plants were transformed by scratching their hypocotyls using 0.5 ml insulin syringe filled with a dense suspension of *A. rhizogenes*. After two weeks of growth on plates under previously described conditions, hairy root formation was observed, and expression of transformed plasmids were checked via optical tYFP-NLS marker. Transformed plants were transferred to sterile 580 ml glasses containing 80 g 3:2 clay:vermiculite mixture and 50 ml ½ strength B&D media inoculated with *M. loti* at an optical density at λ=600 (OD_600_) of 0.01. Plants were grown for four more weeks at long day conditions (16 h light, 21 °C / 8 h dark, 17 °C), 300 µM/s/m^2^ white light before phenotypic evaluation.

### Grafting experiments

Grafting of Lotus seedlings followed a previously described protocol (Sexauer et al., 2023), with minor modifications. Plants were treated and germinated as described above. After germination plants were transferred to ½ –strength B&D / 1 mM KNO_3_ / 1% (w/v) phyto agar medium. Plates were kept three days in long day conditions. For graftings, seedlings were cut near the middle of the hypocotyl and immediately submerged in water. New shoots were transplanted onto root stocks and arrested using silicone tubing (∅∥ 0.5 mm, ∼3 mm long). Grafted plants were transferred to fresh medium, then kept at 26 °C 22 h light for five days to enable graft site regeneration. Afterwards grafted plants were incubated for five more days at long day conditions (16 h light, 21 °C / 8 h dark, 17 °C) before graft success was determined. Successfully grafted plants were infected with *M. loti* and grown for three more weeks at stated conditions before nodulation was evaluated.

### Acetylene reduction assay

For determining nitrogenase activity, two 21 days old plants, or five detached nodules (Fig. 2e) were transferred from petri dishes to 5.3 ml test tubes and the tubes sealed with a gas-tight rubber cap. From a gas sampling bag filled with acetylene (1.15 % in air; All-in-Gas GmbH, Starnberg, Germany) 1 ml was injected by piercing the rubber sealing with a gastight syringe. After 3 h of incubation, 0.5 ml of the headspace was sampled with the gastight syringe and used for gas chromatographic analysis.

For the quantitative analysis a gas chromatograph (Agilent 7890A; Santa Clara, CA, USA) equipped with a flame ionization and split/splitless injector (UNIS 500 S/SL; at 175 °C, split ratio 10:1; Joint Analytical Systems, Moers, Germany) was used. Oven and column (CarboBOND, 25 m x 530 µm x 10 µm; Agilent, Santa Clara, CA, USA) temperature was held constant at 75 °C for 3.5 min with H_2_ as carrier gas (constant flow at 8 ml/min). Acetylene and ethylene contents were quantified by manually integrating peak areas. Nitrogen fixation (*nf*) was defined as:

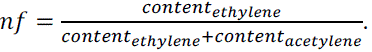

Nitrogen fixation per nodule (*nf/nodule)* was calculated as follows:

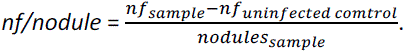

For the normalized fixation per nodule, the average fixation per nodule of wild-type roots was set to 1 for a respective light condition. This was done separately for each replicate of the respective experiment.

### RNA extraction and expression analysis

For total RNA extractions, whole roots or detached nodules were shock frozen in liquid nitrogen and pulverized using stainless steel beads and a bead mill. Total RNA was extracted by a modified Lithium Chloride-Trizol protocol: Crushed tissue was thawed in 400 µl lysis buffer (100 mM Tris-HCl pH 7.5, 0.5 M LiCl, 10 mM EDTA pH 8.0, 1% LiDS, 5 mM DTT) and centrifuged (21,000 × *g*, 10 min, 4 °C). The supernatant was mixed with 600 µl of homemade trizol solution (38% saturated phenol, 0.8 M guanidine thiocyanate, 0.4 M ammonium thiocyanate, 0.1 M sodium acetate pH 5.6, 5% glycerol) and incubated for 10 min at room temperature. Phase separation was achieved by addition of 500 µl chloroform followed by centrifugation (16,000 × *g*, 10 min, 4 °C). The aqueous phase was recovered, mixed vigorously with 500 µl chloroform, and phases were separated again by centrifugation under the same conditions. Subsequently, 500 µl of the aqueous phase were mixed sequentially with 50 µl of 3 M sodium acetate (pH 5.6), 50 µl of 1 M acetic acid, and 1.3 ml of absolute ethanol.

RNA was precipitated by incubation at −20 °C for 22 h, followed by centrifugation at 21,000 x *g* for 1 h at 4°C. The supernatant was discarded, and the pellet was washed twice with ice-cold 80% ethanol and once with 100% ethanol before air-drying at room temperature. RNA was resuspended in 30 µl DEPC-treated water, and concentrations were determined using a NanoDrop spectrophotometer (Thermo Fisher Scientific). Genomic DNA was removed by treatment with DNase I (Thermo Fisher Scientific) according to the manufacturer’s instructions.

cDNA was synthesized from 500 ng of total RNA using RevertAid reverse transcriptase (Thermo Fisher Scientific) with a pulsed thermocycling protocol. Briefly, RNA and oligo(dT) primers (2 µM final concentration) were denatured at 65 °C for 5 min. The remaining reaction components were added, and the mixture was incubated at 16°C for 30 min, followed by 60 cycles of 30°C for 30 s, 42°C for 30 s, and 50 °C for 1 s. The enzyme was inactivated at 85 °C for 5 min.

Quantitative RT-PCR was performed using SensiFAST SYBR No-ROX master mix (Bioline) in 10 µl reactions with a final primer concentration of 500 nM. Reactions were run on a CFX Opus real-time PCR system (Bio-Rad). Expression levels were normalized to two reference genes, *ATP SYNTHASE 2* (LotjaGi5g1v0332700) and *PROTEIN PHOSPHATASE 2A* (LotjaGi2g1v0210500). Primer sequences are listed in Supplementary Table 4. Baseline correction and PCR efficiency calculations were performed using LinRegPCR, relative transcript levels were calculated based on N0 values (Ruijter et al., 2009).

### Microscopy and image acquisition

Symbiotic phenotypes were monitored, and photographs taken using a Leica M205 FCA stereomicroscope equipped with a K7 camera module. Photographs of whole plant phenotypes were taken using an Olympus OMD EM-1 Mark II camera and 17 mm M.Zuiko lens. Semi-thin nodule sections were generated by hand sectioning nodulated root fragments embedded in 6% Agarose blocks.

### Data analysis and graphical representation

Data analysis and generation of graphical representation made use of R 4.3.1 (Team, 2023) using the libraries *multcomp* (Bretz et al., 2016) and *ggplot2* (Wickham, 2016). Boxplot center lines show the medians, outer box limits indicate the 25^th^ and 75^th^ percentiles. Whiskers extend 1.5 times the interquartile range from the 25^th^ and 75^th^ percentiles, or the last data point. Data points are represented as circles. Dotplots center lines indicate the average value of all data points. All statistical tests used were two-sided. For pairwise comparisons, we used t-tests, for multi-comparisons ANOVA and post-hoc Tukey-HSD testing. ANOVA or t-test analysis results, biological replicate numbers and individual datapoint values are listed in the Source Data file.

## Acknowledgements

We thank Jutta Winkler-Steinbeck and the gardening team of the Botanical Garden for dedicated plant care, Christine Gernert for genotyping of LORE1 insertion lines and plant work support. Further we thank Michael Dienesch for technical support with light regime setups. We thank Christoph Weiste for providing SRDX base vectors and Andreas Hiltbrunner for providing *hy5-2* mutant seeds. We also thank Angela Fischer for assistance with graphical representations and photography as well as proofreading of the manuscript.

We apologize to authors whose work could not be cited due to space limitations.

## Author contributions

M.Se., M.A., M.R., S.B., J.R., M.St. and A.L., performed experiments and analyzed data; M.Se., M.A., and K.M. conceived and designed research; K.M. and M.Se. wrote the paper; M.A. and M.R. proofread and corrected the paper.

