## Supplemental material for "Global integrators of light signalling and nutrient homeostasis safeguard symbiosis under light stress"

### Supplementary Information

#### Supplementary Tables

Supplementary Table 1: Primers used for genotyping

| Target | Orientation | Sequence |
|---|---|---|
| *bbx21-1* | forward | GGCTTCAGTATTCTGCACCGCCGA |
| *bbx21-1* | reverse | GGTCTTCAACTTGCCAACCAGGGA |
| *bbx21-2* | forward | GGCTTCAGTATTCTGCACCGCCGA |
| *bbx21-2* | reverse | TGCCATGGTTACCAAGCTTCCTCCA |
| *hy5-2* | forward | TGCCCATTTGTCAGATGAGGATTTTGC |
| *hy5-2* | reverse | TTGGCCTAACAGCCAACGGCGATA |
| LORE1 insertion | reverse | CCATGGCGGTTCCGTGAATCTTAGG |

Supplementary Table 2: Plasmids used for transformation

| Name | Purpose | Features | Origin |
|---|---|---|---|
| *HY5-GFP* | Complementation | *pUBI1:HY5-GFP; pDR5:NLS-tYFP* | This study |
| *HY5-SRDX* | Dominant negative *HY5* | *pUBI1:HY5-SRDX; pDR5:NLS-tYFP* | This study |
| *NLS-GFP* | Empty vector control | *pUBI1:NLS-GFP; pDR5:NLS-tYFP* | This study |

Supplementary Table 3: Primers used for cloning

| Target | Orientation | Sequence |
|---|---|---|
| *HY5* CDS lvl 0 | forward | aGGTCTCaAGGTATGGATCAGCATGGTGGAGT |
| *HY5* CDS lvl 0 | reverse | GGTCTCaCACCTTCAGCATTATTGGTACCAC |

Supplementary Table 4: Primers used for qPCR

| Target | Orientation | Purpose | AT | Sequence |
|---|---|---|---|---|
| *ATPs* | forward | qPCR | 60 | CAATGTCGCCAAGGCCCATGGTG |
| *ATPs* | reverse | qPCR | 60 | AACACCACTCTCGATCATTTCTCTG |
| *PP2A* | forward | qPCR | 68 | GTAAATGCGTCTAAAGATAGGGTCC |
| *PP2A* | reverse | qPCR | 68 | ACTAGACTGTAGTGCTTGAGAGGC |
| *TML* | forward | qPCR | 60 | GCCAACAATTGCCTGAAACCAGATG |
| *TML* | reverse | qPCR | 60 | CTTTATGGTGTTTCTCTCTATGAATGCTG |
| *HY5* | forward | qPCR | 62 | TTGCAACTGGGTCGCATGAAAG |
| *HY5* | reverse | qPCR | 62 | AGCACACTCATGGCAAGCAAC |
| *BBX21* | forward | qPCR | 62 | ACCATGCCAACAAACTCGCC |
| *BBX21* | reverse | qPCR | 62 | TCACACCAGTGAGAAGGAACCG |
| *NIN* | forward | qPCR | 58 | GCCATCAAGGTATATGACGAG |
| *NIN* | reverse | qPCR | 58 | AGGAGCCCAAGTGAGTGCTA |
| *ERN1* | forward | qPCR | 58 | CCTACACTCCTCCCTCTCAAG |
| *ERN1* | reverse | qPCR | 58 | CCACCCTTGTGCTCATTGTTCTG |
| *NSP2* | forward | qPCR | 64 | GGAGGAGCTGGGTAGTAATAAG |
| *NSP2* | reverse | qPCR | 64 | GAGATCTGAAGCGATTTAACAGC |
| *NAC094* | forward | qPCR | 60 | TCAGCCTCAGCCTCAGATATTCC |
| *NAC094* | reverse | qPCR | 60 | ACTTTCCCACTGATTAAGCTGCAC |
| *PR8* | forward | qPCR | 62 | TGGCATTGCTGTCTACTGGG |
| *PR8* | reverse | qPCR | 62 | TTTGCAAGCTGTGTGGCTTC |
| *PR10.2* | forward | qPCR | 55 | CCCAAAGGTTATTCCTGTTTTCA |
| *PR10.2* | reverse | qPCR | 55 | ACGGTTCCGACGGCAAT |

#### Supplementary Figures

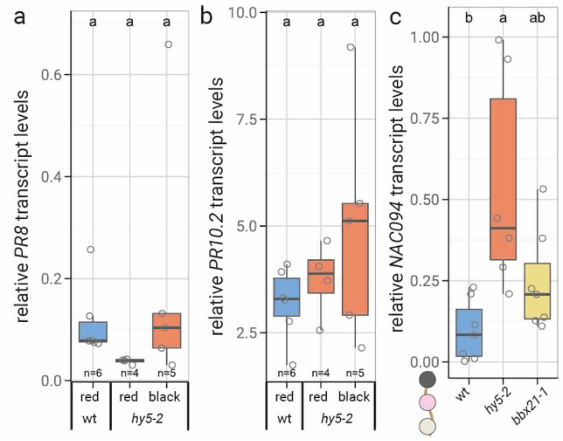


**Supplementary Fig. 1. *hy5-2* nodules accumulate the *NAC094* senescence marker transcripts, but *PR8* and *PR10* defense marker levels are wild type like.** **a**-**b**, *PR8* (**a**), *PR10.2* (**b**) and *NAC094* (**c**) transcript levels in nodules three weeks post inoculation with *M. loti*. **a**-**b**, Pools of five nodules each were harvested for analysis, where red = mature nodules without discoloration and black = nodules showing dark discoloration. **c,** All nodules of two plants per replicate were analyzed, independent of visual appearance. Transcript levels were quantified *via* qPCR relative to two housekeeping genes (*ATPs* and *PP2A*). Plants were evaluated after three weeks of sterile culture. Comparisons used Analysis of Variance (ANOVA) and *post-hoc* Tukey testing (p≤0.05), with distinct letters indicating significant differences p≤0.05 (**b**-**d**) or Students' t-test (**a**). Boxplot central line shows median value, box limits indicate the 25$^{th}$ and 75$^{th}$ percentiles. Whiskers extend 1.5 times the interquartile range or to the last datapoint. Individual data points are represented by dots.

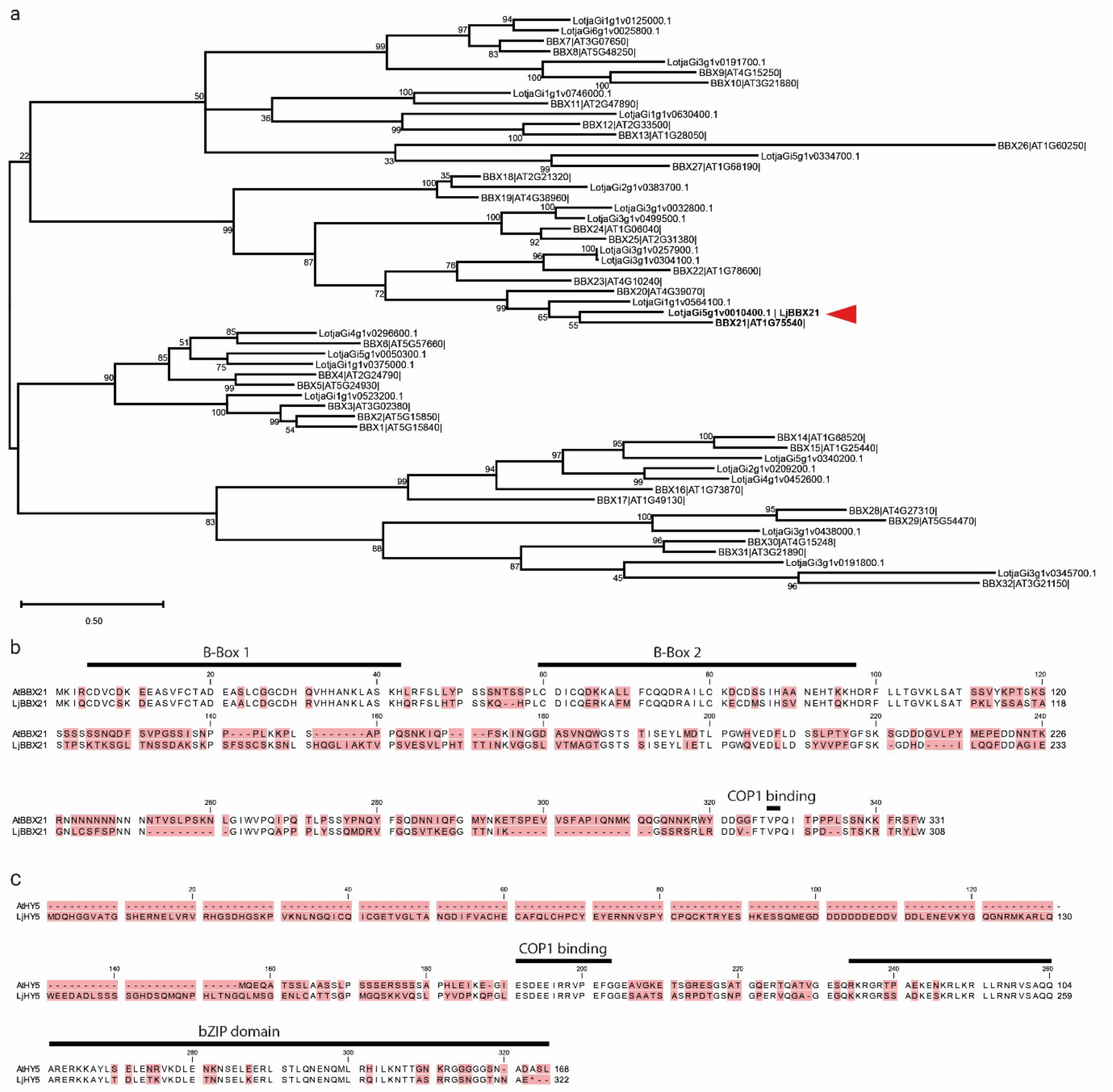


**Supplementary Fig. 2. Phylogenetic analysis identifies a putative *At*BBX21 orthologue in *Lotus japonicus*. a,** Comparative sequence analysis predicting phylogenetic relations of 32 *A. thaliana* and 23 *L. japonicus* putative BBX proteins. Analysis was based on full-length amino acid sequences. Sequences were aligned using ClustalW algorithm (Thompson et al., 1994). The phylogeny was inferred using the Maximum Likelihood method and Jones-Taylor-Thornton model of amino acid substitutions (Jones et al., 1992). The tree with the highest log likelihood value is shown. Numbers indicate the percentage of replicate trees (total: 500) in which the respecticve taxa clustered together (Felsenstein, 1985). The initial tree for the heuristic search was selected by choosing the tree with the superior log likelihood between a Neighbor-Joining (NJ) tree (Saitou and Nei, 1987) and a Maximum Parsimony (MP) tree. The NJ tree was generated using a matrix of pairwise distances computed using the Jones-Taylor-Thornton model (Jones et al., 1992). The MP tree had the shortest length among 10 MP tree searches; each performed with a randomly generated starting tree. Evolutionary analyses were conducted in MEGA12 (Kumar et al., 2024). *Lj*BBX21 is marked by a red arrow. **b-c,** Alignments of *At*BBX21 and *Lj*BBX21 (**b**) or *At*HY5 and *Lj*HY5 (**c**). *Lj*, *Lotus japonicus*; *At*, *Arabidopsis thaliana*. **b-c,** Alignments were generated using CLC Main Workbench 25 (Qiagen) software. Red background indicates non-conserved residues.

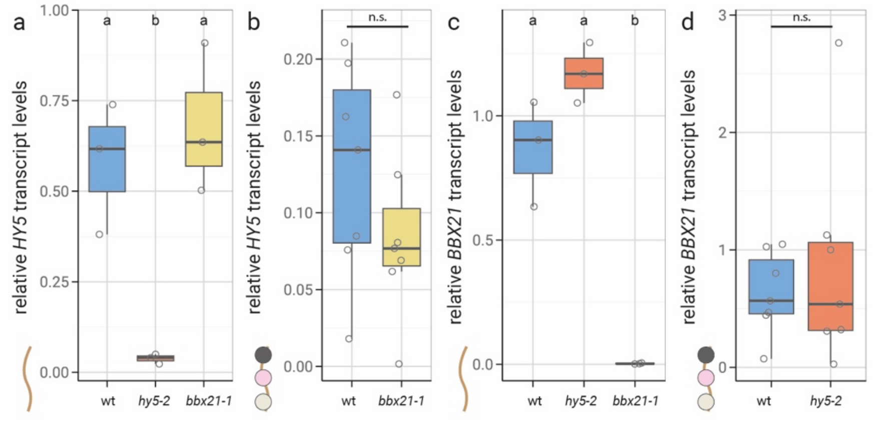


**Supplementary Fig. 3. *HY5* and *BBX21* show no mutual transcriptional dependencies. a-d,** Relative *HY5* (**a**,**b**) and *BBX21* (**c**,**d**) levels in Lotus roots (**a**,**c**) or detached nodules (**b**,**d**). Transcript levels measured via qPCR, relative to the housekeeping genes *ATPs* and *PP2A.* Tissue harvested 3 weeks post infection with *M. loti*. Comparisons used Analysis of Variance (ANOVA) and *post-hoc* Tukey testing ($p \leq 0.05$) (**a**,**c**), with distinct letters indicating significant differences $p \leq 0.05$, or Students' t-test (**b**,**d**). Boxplot central line shows median value, box limits indicate the 25$^{th}$ and 75$^{th}$ percentiles. Whiskers extend 1.5 times the interquartile range or to the last datapoint. Individual data points are represented by dots.

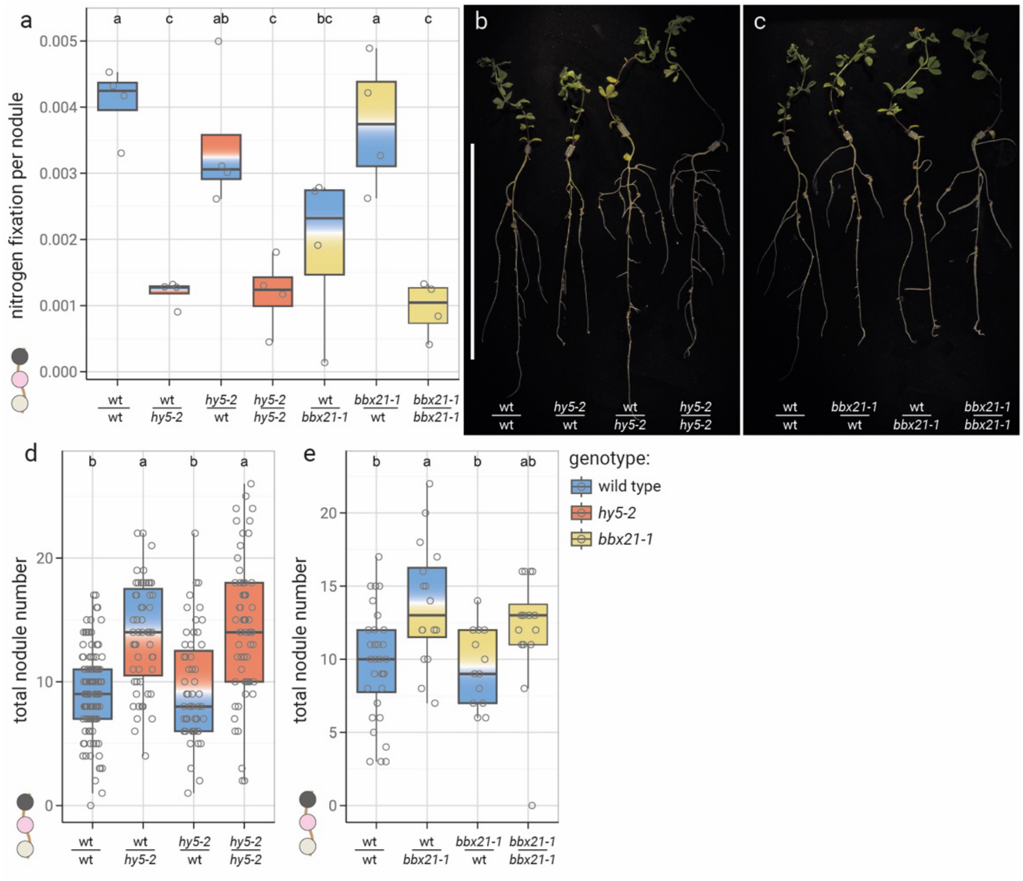


**Supplementary Fig. 4. Root *HY5* and *BBX21* ensure nitrogen fixation and balance nodule numbers. a-e**, Nitrogen fixation per nodule (**a**), whole plant growth phenotypes (**b**,**c**) and total nodule number (**d**,**e**) of grafts between *L. japonicus* Gifu wild type (wt) and *hy5-2* (**a**,**b**,**d**) or *bbx21-1* (**a**,**c**,**e**) plants. Scale bar equals 10 cm. Plants evaluated 3 weeks after inoculation with *M. loti*. Comparisons used Analysis of Variance (ANOVA) and *post-hoc* Tukey testing (p≤0.05), with distinct letters indicating significant differences p≤0.05. Boxplot central line shows median value, box limits indicate the 25th and 75th percentiles. Whiskers extend 1.5 times the interquartile range or to the last datapoint. Individual data points are represented by dots.

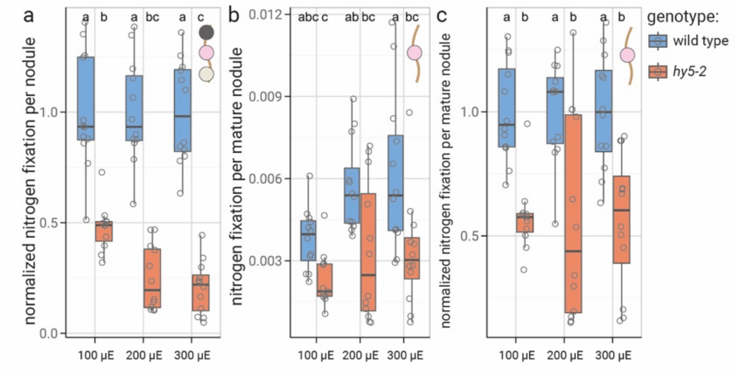


**Supplementary Fig. 5. Light intensity affects nitrogen fixation in *L. japonicus* nodules. a**-**c**, Normalized nitrogen fixation per nodule (all nodules considered; **a**), nitrogen fixation per mature nodule (**b**), and normalized nitrogen fixation per mature nodule (**c**) of wild type (wt) and *hy5-2* plants. Nitrogen fixation divided by number of total nodules, including non-fixing immature and senescent nodules (**a**) or divided by number of mature nodules (**b**, **c**). **a**, **c**, Nitrogen fixation per mature / total nodule normalized by setting average fixation per nodule of wild-type plants to 1. Plants evaluated 3 weeks post infection with *M. loti.* Plants were grown at indicated light intensities of white light, roots were covered from direct light exposure. Comparisons used Analysis of Variance (ANOVA) and *post-hoc* Tukey testing (p≤0.05), with distinct letters indicating significant differences p≤0.05. Boxplot central line shows median value, box limits indicate the 25$^{th}$ and 75$^{th}$ percentiles. Whiskers extend 1.5 times the interquartile range or to the last datapoint. Individual data points are represented by dots.

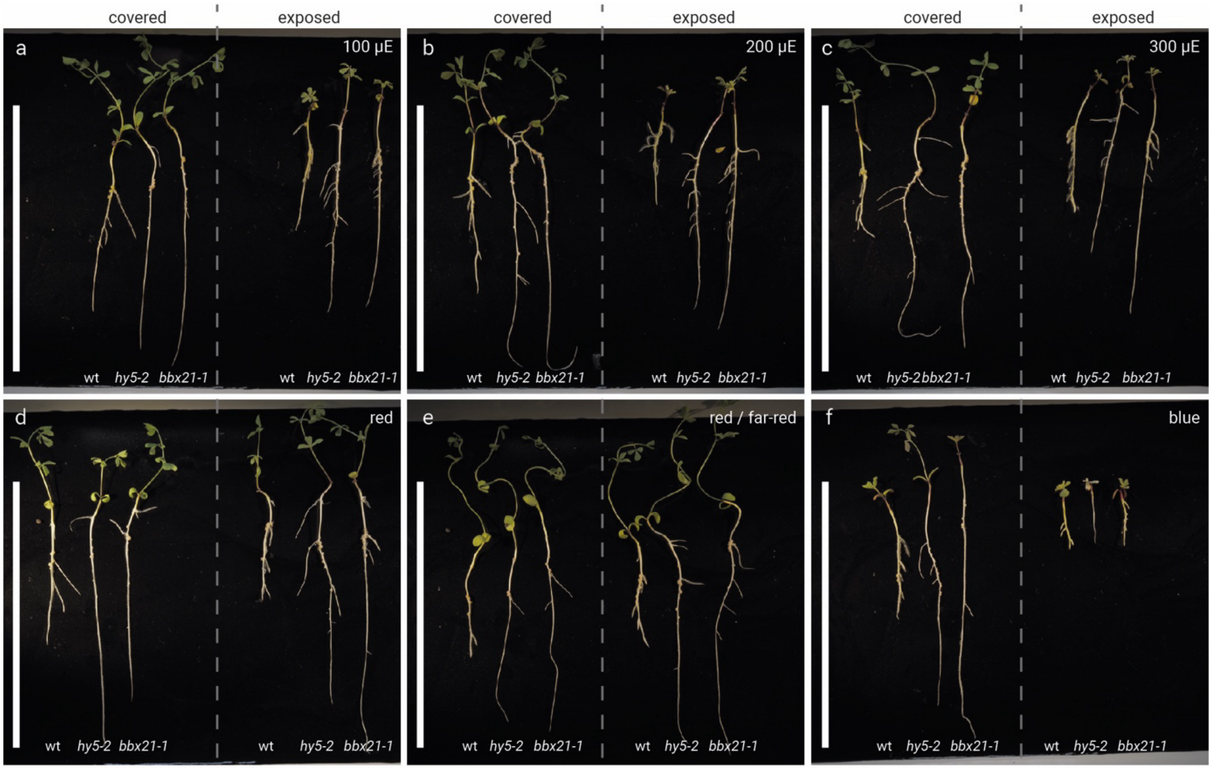


**Supplementary Fig. 6. Light intensity and color influence *L. japonicus* (Lotus) growth and performance. a**-**f**, Whole plant growth phenotypes of Lotus wild type, *hy5-2* and *bbx21-1* plants grown at indicated light conditions. **a**-**c**, Plants were grown under white light at intensity levels of 100 µE (**a**), 200 µE (**b**) or 300 µE (**c**). **d**-**f**, Plants were grown under red ($\lambda$=660 nm) light at 200 µE (**d**), under red as well as far-red ($\lambda$=730 nm) light at 200 µE (**e**) and under blue ($\lambda$=450 nm) light at 200 µE (**f**). Plants on the left side of each picture were grown with roots shielded from direct light exposure, while plants on the right had their roots exposed to the respective light conditions. Plants were infected with *M. loti* and grown for three weeks under the indicated conditions. Scale bars equal 10 cm.

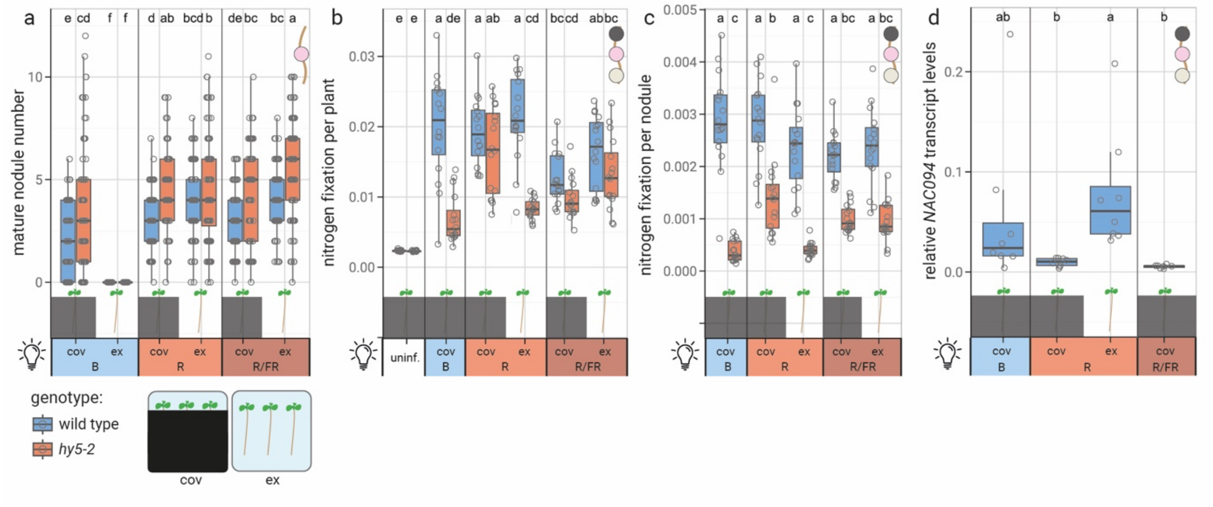


**Supplementary Fig. 7. Red light exposure of roots induces nodule senescence via *NAC094* induction. a-c**, Mature nodule number (no discoloration) (**a**), nitrogen fixation per plant (**b**) and nitrogen fixation per nodule (**c**) of *L. japonicus* Gifu wildtype (wt), *hy5-2* and *bbx21-1* plants grown under indicated light conditions. **d**, Relative *NAC094* transcript levels in wt nodules (data are also shown in Fig. 5d at a different scale). **e**-**h,** Total nodule number (**e**), discolored nodule number (**f**), normalized nitrogen fixation per nodule (**g**) and relative *NAC094* transcript levels (**h**) of *L. japonicus* Gifu wild type (wt), *hy5-2* and *bbx21-1* plants grown under indicated light conditions. Plants were evaluated three weeks post inoculation with *M. loti*. R = red light (660 nm); R/FR = red light (660 nm) with far-red supplementation (730 nm); B = blue light (450 nm); cov = roots covered from light exposure, ex = roots exposed to light. Light intensity was 200 µE. Comparisons used Analysis of Variance (ANOVA) and *post-hoc* Tukey testing ($p \leq 0.05$), with distinct letters indicating significant differences $p \leq 0.05$. All data presented in the respective panels was included in the analyses, vertical separators are visual aids only. Boxplot central line shows median value, box limits indicate the 25$^{th}$ and 75$^{th}$ percentiles. Whiskers extend 1.5 times the interquartile range or to the last datapoint. Individual data points are represented by dots.

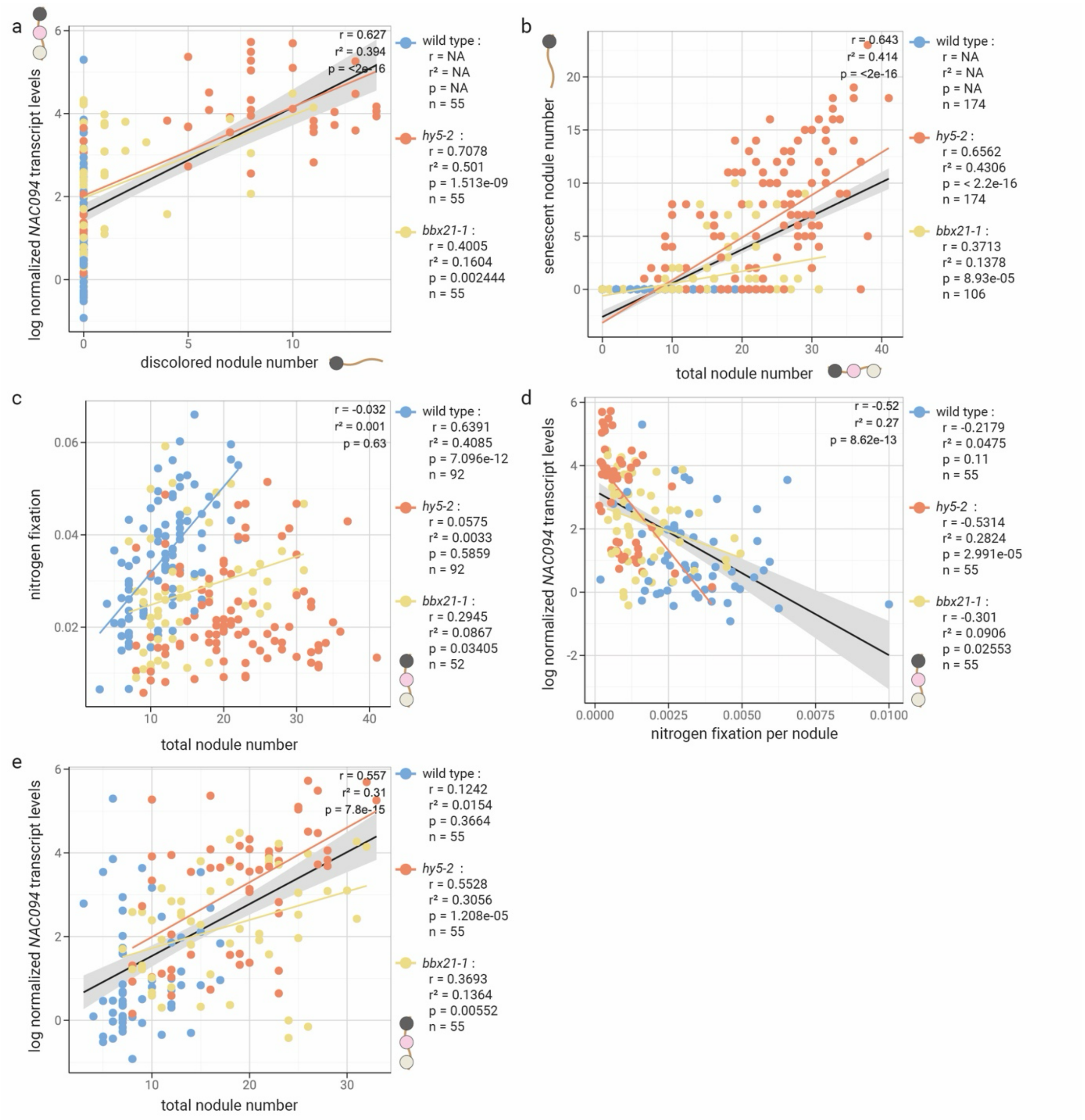


**Supplementary Fig. 8. *NAC094* transcript levels correlate with nodule discoloration and proliferation.**
**a-e**, Correlation plots. Analyses include data shown in **Fig. 2d**, **Supp Fig. 5b**; **Fig. 5d** and **Supp. Fig. 1c**. Pearson correlation analysis was conducted for each dataset as a whole (datapoints of all genotypes were pooled) and for genotype-specific subgroups. Respective results parameters of the genotype specific analyses are displayed next to the graphs. Where a significant correlation ($p<0.05$) was observed, trendlines are displayed. For genotype specific data, the color code is as indicated to the right of the graphs. Where datapoints of all genotypes were pooled, trendlines are in black, with grey areas indicating 95% variance limits. log, natural logarithm.
